# Assessment of coral restoration’s contribution to reef recovery using machine learning

**DOI:** 10.1101/2020.12.10.418715

**Authors:** Gaétan Morand, Simon Dixon, Thomas Le Berre

## Abstract

Coral restoration emerged globally as a form of life support for coral reefs, awaiting urgent mitigation of anthropogenic pressure. Yet its efficiency is difficult to assess, as ambitious transplantation programs handle hundreds of thousands of fragments, with survival rates inherently time-intensive to monitor. Due to limited available data, the influence of most environmental and methodological factors is still unknown.

We therefore propose a new method which leverages machine learning to track each colony’s individual health and growth on a large sample size. This is the first time artificial intelligence techniques were used to monitor coral *at a colony scale*, providing an unprecedented amount of data on coral health and growth. Here we show the influence of genus, depth and initial fragment size, alongside providing an outlook on coral restoration’s efficiency.

We show that among 77,574 fragments, individual survival rate was 31% after 2 years (21% after 4 years), which is much lower than most reported results. In the absence of significant anthropogenic pressure, we showed that there was a depth limit below which *Pocillopora* fragments outperformed *Acropora* fragments, while the opposite was true past this threshold. During the mid-2019 heatwave, our research indicates that *Pocillopora* fragments were 37% more likely to survive than *Acropora* fragments.

Overall, the total amount of live coral steadily increased over time, by more than 3,700 liters a year, as growth compensated for mortality. This supports the use of targeted coral restoration to accelerate reef recovery after mass bleaching events.

## 1 Introduction

Coral reefs are among the most biodiverse ecosystems on earth, yet they are highly threatened by human activity [1], and their rapid degradation is expected within the next 20 years [2, 3]. We are now past the point where we could hope to protect them with the traditional habitat conservation approach; active restoration seems to be essential to their survival, along with a fast mitigation of anthropogenic pressure [4, 5].

Studies relating to the performance of coral restoration have been limited by two factors: the amount of recorded data, and the time and resources needed to interpret it [6]. As a result, coral restoration monitoring programs are often restricted to the short term and bounded to a limited number of observations [7]. Yet scientific uncertainty threatens the quality of management decisions, especially in such a time-critical context [8]. A proper efficacy assessment is vital to a coherent allocation of time and resources.

The increase in frequency of mass bleaching events is threatening reef resilience and its ability to recover [9, 10]. Following the traumatic 1998 mass bleaching event, Reefscapers (formerly Seamarc) started to practice coral restoration, as a recommended way to accelerate reef recovery [11, 12]. The program resulted in a significant increase in hard coral cover and structural reef complexity [13]. It is also a way to engage with the public and educate both local communities and foreign tourists about the fate of coral reefs [14].

Branching species are high rate carbonate producers and they are being replaced by low-relief coral forms, which leads to a decrease in diversity and complexity [15]. To increase natural recovery rate, we attach fragments from the *Acropora* and *Pocillopora* genera onto metal frames, following the direct transplantation best practices [16]. Our objective is to monitor each frame every 6 months, taking pictures from four angles. On average, our 3,368 active coral frames were monitored every 187 days (standard deviation 60 days). This constitutes a significant photographic database, so we constructed a new method to efficiently analyze monitoring data on a large scale. We successfully implemented computer vision algorithms to process monitoring pictures and extract the information they contain.

Artificial intelligence techniques have already been proven to outperform humans in a diversity of settings [17, 18], and they were recently suggested for use in the field of coral conservation [19]. Machine learning was used to the classify coral taxa in survey pictures [20], and even to identify coral species [21]. We analyzed monitoring pictures using convolutional neural networks to locate coral colonies and the metal frames [22, 23]. It enabled us to track each individual colony in a cost effective and time-efficient manner [24], which lead to the generation of a large health and growth database [25]. We could then study the influence of environmental disturbances on mortality rates. The large sample size also made it possible to establish key success factors for direct transplantation.

## 2 Results

### 2.1 Impact of global and local pressure

In order to present results that were independent from local conditions, we first looked at potential local variations. To do this, we split our fragments by geographic areas, which showed a strong disparity in mortality rate (see figure 1).

**Figure 1:**
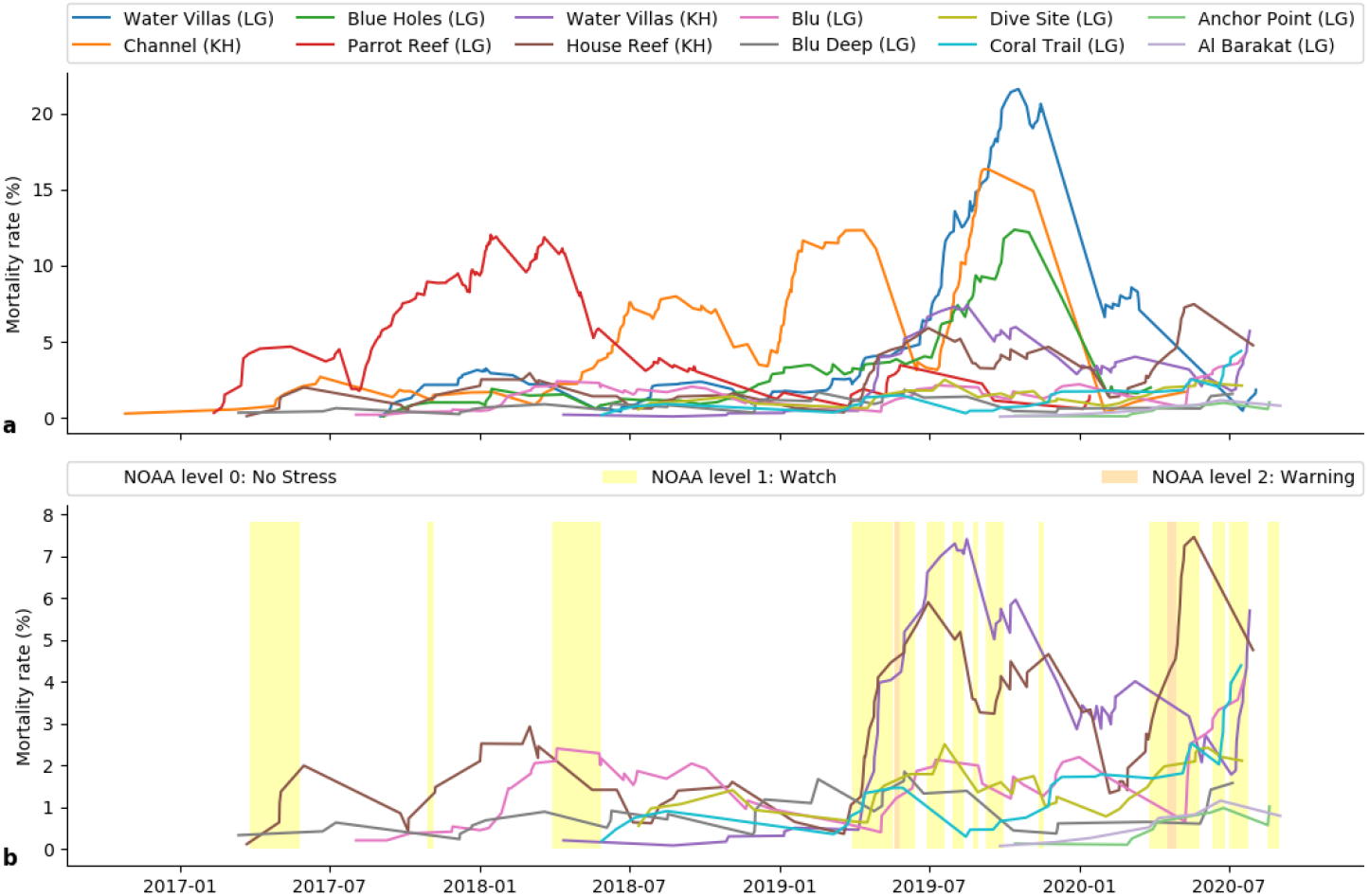
Evolution of the mortality rate between successive monitorings. **a:** All zones with more than 1,000 detected fragments. **b:** Same zones as a, except those with high locally induced mortality [Water Villas (LG), Channel (KH), Blue Holes (LG), Parrot Reef (LG)]. Solid colors show the NOAA bleaching alert level for Kuda Huraa.

Figure 1a highlights significant mortality peaks for four different zones:

- Water Villas (LG): We observed a very localized peak between June 2019 and January 2020. Significant construction work took place in this area at the time, which lead to coral frame damage, high turbidity and increased sedimentation.
- Channel (KH): There are several mortality peaks, which can be attributed to the high sedimentation levels caused by nearby dredging. A land reclaiming project on the neighbour island of Bodu Huraa took place in several phases at this time.
- Blue Holes (LG): These frames were located at the Water Villas (LG) at the time of the peak; see above.
- Parrot Reef (LG): We suspect significant *Drupella* sp. predation until the end of 2018, as the preceding 2016 mass bleaching made the small amount of remaining live coral vulnerable. Parrot Reef used to be a thriving reef before that bleaching event, while most of our other zones are more ecologically isolated, which might explain the localization of *Drupella* predation.

In figure 1b, we removed the previously mentioned zones to have a clear view of those remaining. We noticed two significant peaks at the Water Villas (KH) and the House Reef (KH), both situated in Kuda Huraa. We observed a strong correlation with the NOAA alert level at this location, which suggests that this mortality was caused by prolonged high sea surface temperature, likely caused by climate change [26].

We defined two separate subsets of fragments for the rest of the survival study:

- The disrupted group (N = 54,387 fragments) which includes frames from the 6 zones that suffered high time-specific mortality: Water Villas (LG), Channel (KH), Blue Holes (LG), Parrot Reef (LG), Water Villas (KH), House Reef (KH)
- The control group (N = 21,241 fragments), composed of the frames situated in the rest of the zones with more than 1,000 fragments: Blu (LG), Blu Deep (LG), Dive Site (LG), Coral Trail (LG), Anchor Point (LG), Al Barakat (LG)

Figure 2 shows the average survival rate of our fragments, using the Kaplan-Meier non-parametric curves to take into account the fact that most of our survival times were unknown (fragments are still alive on the reef) [27]. We observed an average 31% survival rate after 2 years (36% for the control group). The average survival rate dropped to 21% after 4 years.

**Figure 2:**
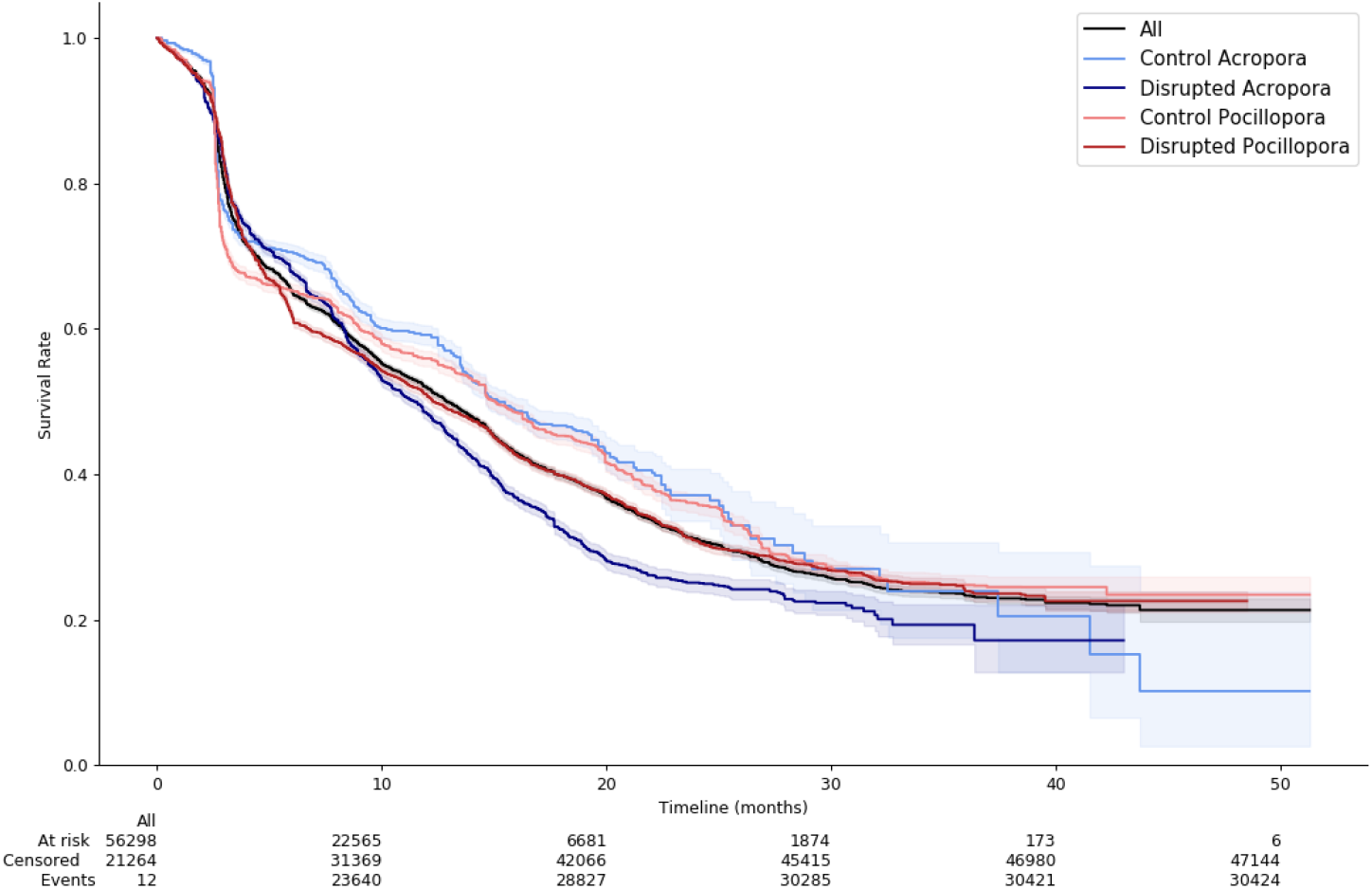
Survival rate of the previously defined four subgroups. The colored area shows the 95% confidence interval. The table shows, for each duration, the number of coral fragments that were still alive (At risk), that were dead (Events), and which survival time was longer but unknown (Censored)

### 2.2 Influence of genus, depth and initial fragment size in normal conditions

After obtaining global survival results, we took advantage of the large number of fragments to analyze the influence of methodological and environmental factors on survival. We singled out genus, initial fragment size, and depth as variables that we could study and that might impact survival rate. We used Cox model [28] to fit regression parameters and establish the influence of these parameters. We also added a binary variable that is equal to 1 when the fragment is on the bottom bar of the frame, in a sandy environment. The Cox model assumes that all fragments have the same mortality rate (function of time), modulated by a hazard ratio: a factor that depends on the covariates but not on time. For the control group, this hazard ratio can be calculated for *Acropora* and *Pocillopora* respectively:

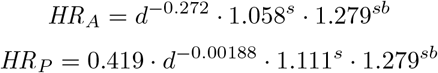

Where *d* represents the depth (m), *s* the initial fragment size (cm), and *sb* the binary variable defining whether the fragment is on the bottom bar in a sandy environment.

Immediately we notice that *Acropora* fragment survival is very dependent on depth, while *Pocillopora* fragment survival is more dependent on fragment size.

Figure 3 shows how the hazard ratio (*ie*. mortality multiplier) changed with regards to depth. It demonstrates that mortality rates were higher at shallower depths, and with larger initial fragments.

**Figure 3:**
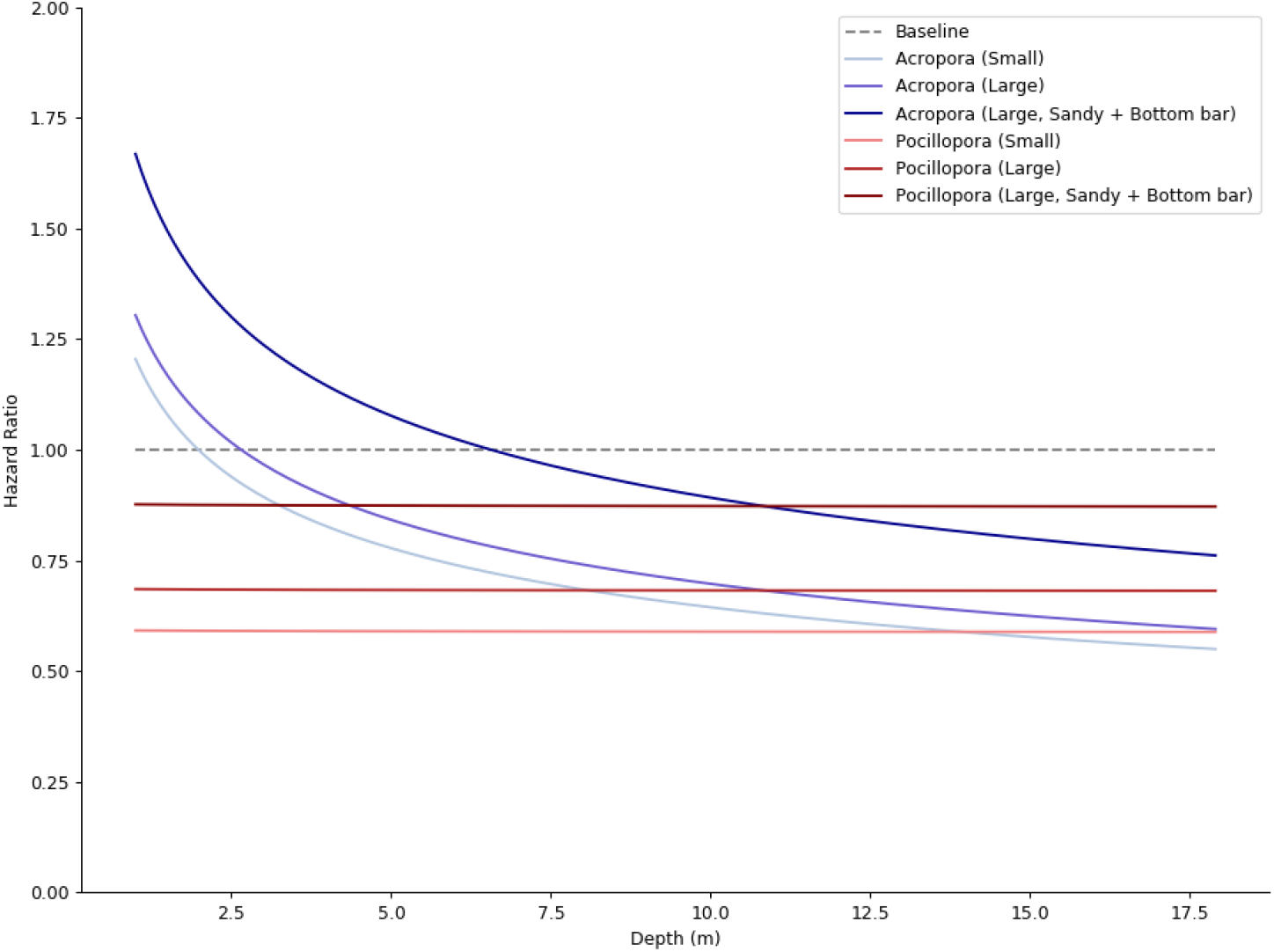
Evolution of the hazard ratio for *Acropora* and *Pocillopora*, for initial sizes representing the first and third quartiles of our data (resp. 3.3 cm and 4.7 cm).

Interestingly for future methodology updates, it is clear that *Pocillopora* fragments performed better than *Acropora* fragments at shallower depths, while *Acropora* slightly had the edge at greater depths. That depth threshold increased as initial fragment size decreases.

We also highlight the higher mortality that occurred on the bottom bars of frames in sandy environments. Indeed, they tended to sink into the sand or be partly covered due to sand movement.

### 2.3 Influence of genus and initial fragment size on vulnerability to heatwaves

Unfortunately, the disparity in the nature and intensity of observed environmental anomalies prevented us from analyzing survival in affected areas as a whole. We focused on the two zones that presumably suffered from the mid-2019 heatwave: Water Villas (KH) and House Reef (KH). We could not include depth in this analysis as it was strongly correlated to many irrelevant factors in such a localized sample. For instance, all the Water Villas (KH) frames are shallow (1-2 m), and almost all of the House Reef (KH) frames are deep (10-11 m).

We introduced a time-varying binary variable that was equal to 1 during the main heatwave event, *ie*. between March 30th, 2019 to September 30th, 2019. We did not take into account the 2020 heatwave as subsequent monitoring data is insufficient as of now. We used Cox model adapted to time-dependent variables to infer the correlation between survival and our covariates [29]. Results must be interpreted with caution as we were not able to check the validity of the proportional hazards hypothesis in this section, nor use the robust inference method.

We show in figure 4 the impact of initial fragment size, genus and disturbance on the survival rate. *Acropora* fragments seemed to be much more vulnerable to the disturbance than *Pocillopora* fragments. We did not find significant correlation between initial fragment size and vulnerability to the disturbance. In accordance with the previous section, we observed a positive correlation between initial fragment size and mortality rate.

**Figure 4:**
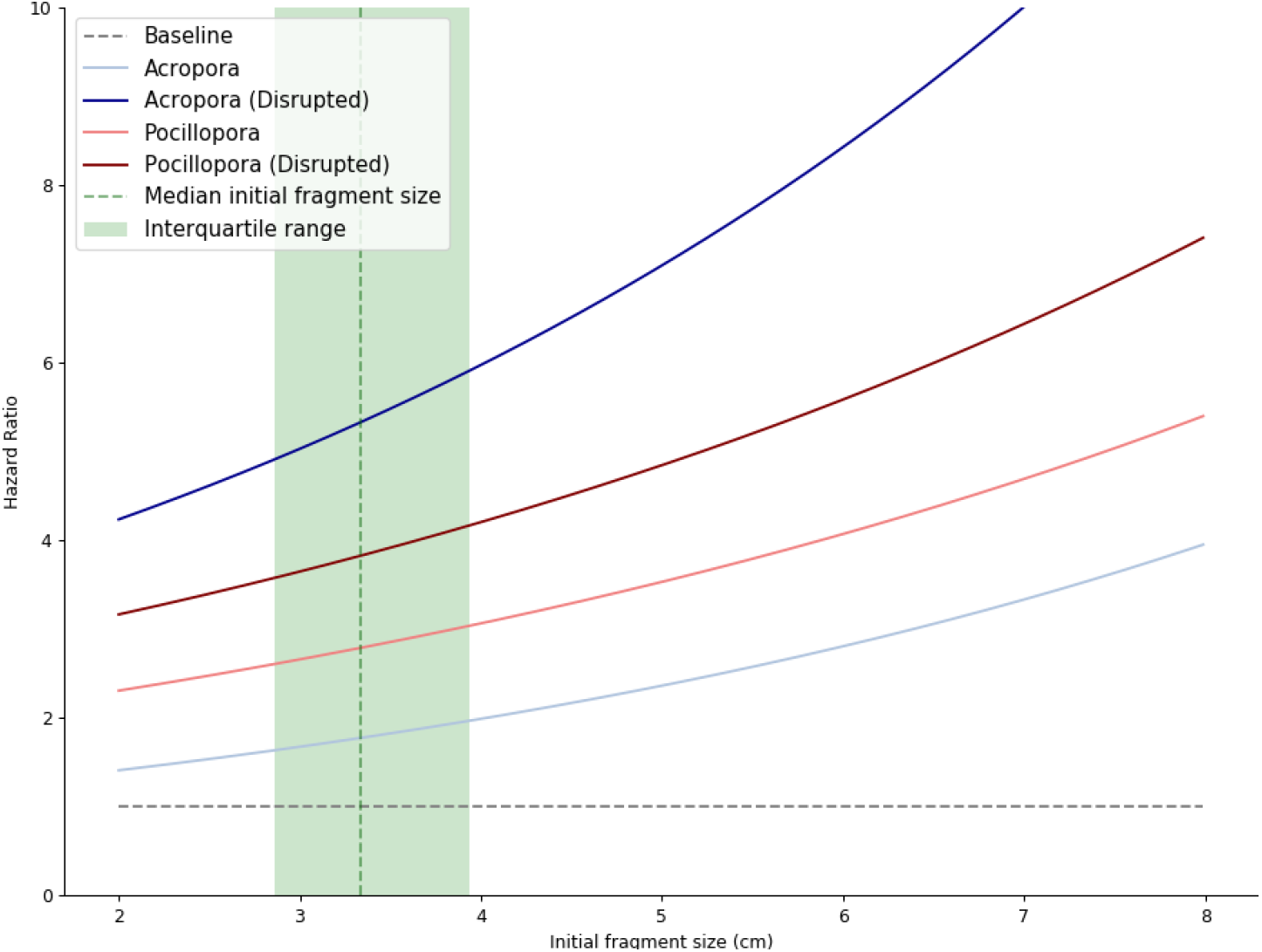
Evolution of the hazard ratio for *Acropora* and *Pocillopora*, with regards to initial fragment size. Local results for the two areas that were presumably affected by the 2019 heatwave: House Reef (KH) and Water Villas (KH). The green dashed line and area show the median and the central 50% of our initial fragment sizes respectively.

In these two geographical zones, this hazard ratio can be calculated for *Acropora* and *Pocillopora* respectively:

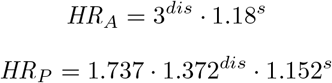

where *s* is the initial fragment size and *dis* the binary variable that is equal to 1 in the presence of a heatwave.

For the median initial fragment size (3.33 cm) and in the absence of disturbance, *Acropora* fragments had a 38% lower mortality rate than *Pocillopora*. During the disturbance, the odds were reversed and *Pocillopora* fragments had a 27% lower mortality rate than *Acropora*. We emphasize that these results are very specific to the location and to the mid-2019 heatwave ; they should not be generalized.

### 2.4 Statistics on deaths and bleaching

In this section we considered fragments from all geographical zones again. Our results show that among the 30,424 recorded dead fragments, 3,066 (10.1%) were detected as dead during later monitoring events, while the rest, 27,358 (89.9%), were never detected again. This latter part is comprised both of fragments falling from the frame while still alive, fragments which died and consequently fell, and fragments that were buried due to sand movement. This percentage is much higher for *Acropora* (13.1%) than for *Pocillopora* (8.2%). This suggests that *Pocillopora* fragments have a higher tendency to fall from the frame while still alive, hence unnecessarily reducing chances of survival.

Among the 600 fragments that were detected as bleached, 358 were monitored again. Among those, we observed a 58% recovery rate for *Acropora* fragments and 61% for *Pocillopora* fragments. The rest of the bleached fragments were observed more recently and were not monitored since.

Bleaching occurred within 15 days following transplantation in 17.2% of cases, which was most likely a result of transplantation stress. 95% of these fragments recovered.

For the remainder of the fragments, bleaching occurred due to environmental stress. Indeed, 82.9% of the fragments in this group were located in one of the four zones affected by local pressure: Water Villas (LG), Channel (KH), Blue Holes (LG) and Parrot Reef (KH), which make up 40.7% of the total fragment number. But only 5.8% of them were located in the areas presumably affected by heatwaves: Water Villas (KH) and House Reef (KH), which represent 25.2% of the total fragment amount.

### 2.5 Growth rates

We analyzed coral growth by measuring the evolution of a theoretical radial length for each live colony (see methods section). This measurement is quite different from linear branch growth, as it averages the 3D growth of the colony. Overall, we observed a 1.05 cm/year median growth rate for *Acropora* fragments and 1.33 cm/year for *Pocillopora* (respective interquartile ranges: 1.12 cm/year and 0.98 cm/year).

Each data point (N=32,265) represents a single fragment. The growth of each fragment is the average of all size variations, measured between any two monitoring dates (not only consecutive ones). This enabled us to smooth out statistical errors. We excluded such size differences when there was less than 5 months between the two monitoring sessions; in this case the detection error might be more significant than the growth we are measuring.

We observed a strong disparity of growth rates among geographical zones and fragment genera. Figure 5 shows the distribution of growth rates. For each category, the black line illustrates the full range of data points, while the colored areas show their distribution. These outliers are mostly consequences of sporadic detection mistakes by the machine learning models (see methods section).

**Figure 5:**
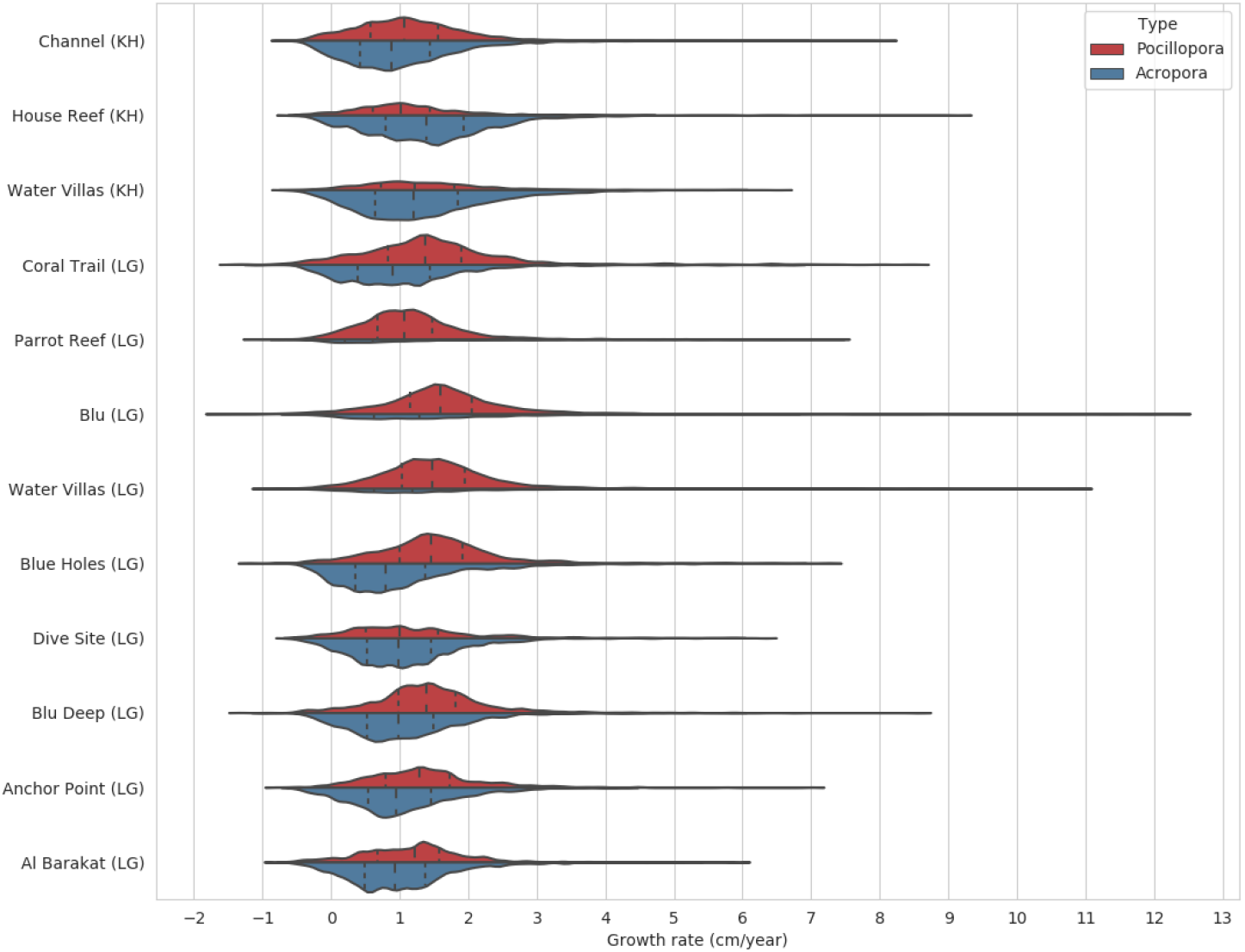
For each geographical zone, distribution of growth rates for *Acropora* and *Pocillopora* fragments. Long dashes depict median, short dashes depict first and third quartiles. Width is proportional to the number of observations.

In the most successful areas, we observed a 1.38 cm/year median growth rate for *Acropora* and 1.58 cm/year for *Pocillopora* (House Reef (KH) and Blu (LG) respectively). The poorest performing areas saw a 0.61 cm/year median growth for *Acropora* and 0.99 cm/year for *Pocillopora* (in Parrot Reef (LG) and Dive Site (LG) respectively).

### 2.6 Total volume

Quantifying shelter is an important step towards assessing the restored reef’s capacity to support biodiversity [30]. We infered the shelter volume (*ie*. the convex hull volume) of each colony for every monitoring set (see methods). We used a conservative estimate as we did not account for growth in-between measurements and we assumed the volume at any time is the volume at the last monitoring date. We could then classify that volume into four mutually exclusive categories at any point in time:

- **Live coral, initial transplants**. When a fragment is first detected, the recorded volume is the volume that was moved from the donor colony to the coral frame.
- **Live coral, organic growth**. In the subsequent monitoring pictures, the difference between the current colony’s volume and the initial transplant’s volume was generated by organic growth.
- **Dead coral, on frame**. Some of the colonies that die stay attached to the frame. We choose to measure that volume as well as it still provides a habitat to sea life.
- **Dead coral, fallen**. Some of the colonies fall from the frame, either before or after dying. They participate in the formation of bottom sediments and contribute to the benthic habitat.

Figure 6 shows the evolution of these four categories through time, for *Acropora* (6a) and *Pocillopora* (6b). It illustrates that the volume of live coral steadily increased. We calculated an approximate slope for this increase from January 2019: 981 liters/year for *Acropora* and 2,748 liters/year for *Pocillopora*.

**Figure 6:**
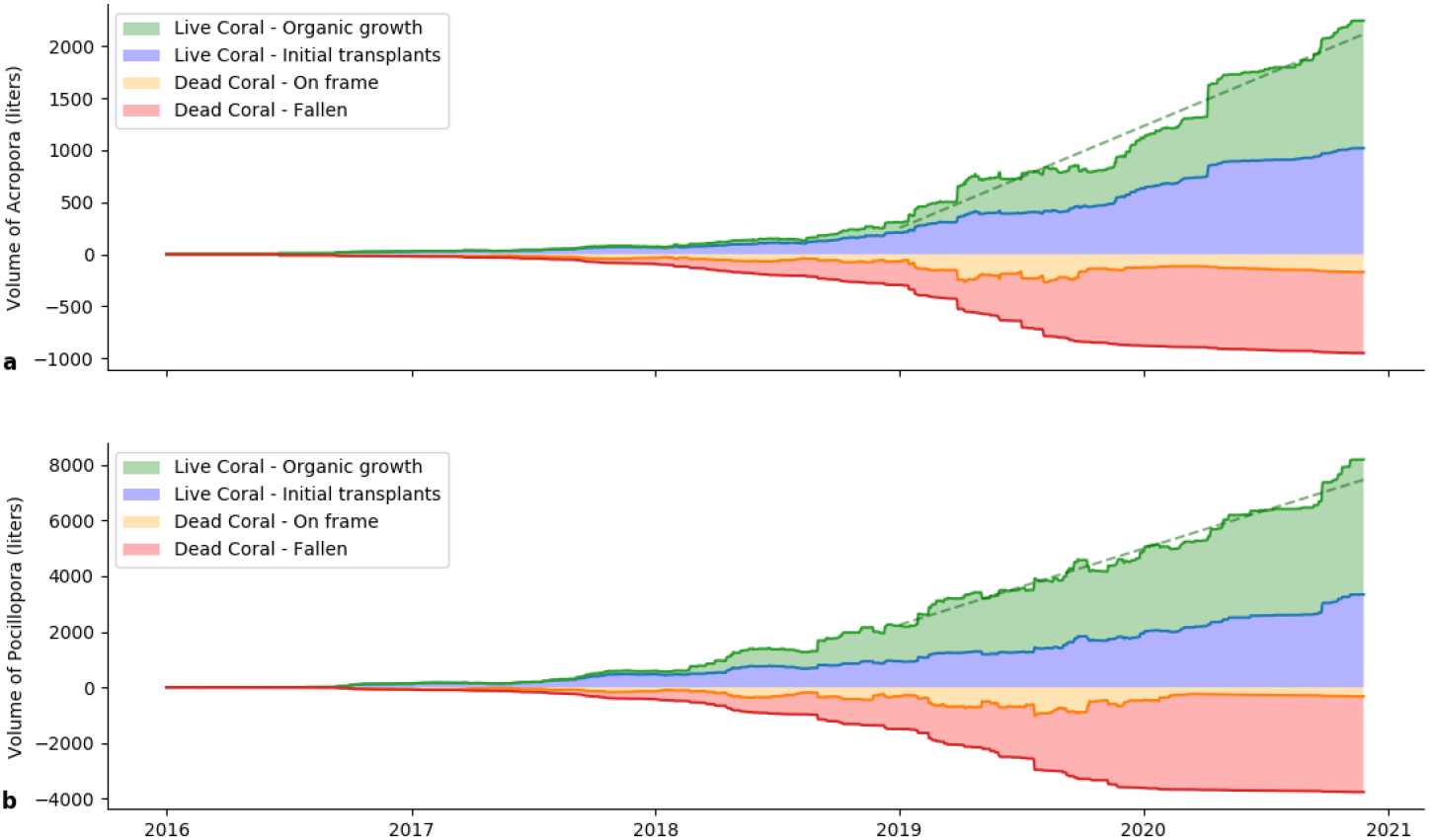
Evolution of the volume of live coral (positive) and dead coral (negative). The green and red lines show the total amount of live and dead coral respectively. The four colored areas represent the four categories previously described. Dashed lines show the trend lines for live coral since January 2019.

## 3 Discussion

### Time frame

The time period analyzed in this study does not include mass bleaching events, which threaten the survival of the vast majority of both wild and transplanted coral colonies [31]. Coral restoration is a tool to help the recovery of coral reefs after such events, but it can not aim to prevent or fix such mass mortality [5].

The method we described is designed to analyze coral restoration’s ability to facilitate coral recovery. We focus on providing colony-level observations on a large number of fragments in their first 3-4 years after transplantation. This method is not adapted to a longer time frame, as colonies then tend to merge together; it makes it impossible to keep tracking them individually. Other methods (which might also include machine learning) are more adapted [24], and would enable studying success on the longer term, a beneficial complement to this study.

### Survival

In this study we provide a multitude of different indicators to assess the outcome of the coral restoration program on a colony level. The survival rate we report (31% after 2 years) is much lower than the usually reported 60-70% [6]. But thorough review points out frequent exaggerated survival results, through cherry-picking of successful metrics, non-representative fragment subsets, as well as accidental mortality underreporting [32]. The exhaustiveness of our monitoring might explain our higher than average mortality rate, although local conditions specific to our protocol or to the Maldives cannot be excluded.

Furthermore, most studies on transplanted fragment survival encompass a limited geographical area, which makes the results specific to that area [33, 34]. We overcome this caveat by including fragments transplanted according to the same protocol in 12 different geographical areas, in a diversity of substrates, depths, and water circulation conditions. Mortality results in section 2.1 highlight the disparity of mortality rates among different geographical areas, which confirms the need for a large sample size distributed over as many different areas as possible. Most existing survival and growth studies also rely on a limited number of fragments (a few hundred at best) [35, 36], either because of the experiment’s small scale, or because only a small subset was monitored due to time constraints. Nothing guarantees the representativeness of such subsets, and the low numbers prevent any reliable analysis of influential factors.

As we produced data that is more detailed than what usual survey time constraints would allow, we introduced new metrics to take advantage of the additional information. For example, we proposed a live coral volume metric, a reliable measurement of the habitat that restoration provides to marine life; it depicts the real environment in its three dimensional structural complexity [37]. For transparency and standardization purposes, we publish all our raw and processed data in an open database [25].

### Implications for the future

The results we provide need to be used with caution, as they highly depend on the accuracy of all our method blocks (see methods section). Yet small confidence intervals and large sample size in section 2.2 enable us to confidently recommend transplanting an assemblage of *Pocillopora* and *Acropora*, as they have complementary roles. *Pocillopora* seems to be more resistant to environmental perturbations, while *Acropora* provides more species to contribute to genetic diversity [38]. This mix should consist in a majority of *Pocillopora* at shallower depths (less than 5m), as this genus will have significantly higher chances of survival. This corroborates previous studies that showed depth could act as a refuge for *Acropora* during bleaching events [39].

It is critical to study the effects of weather patterns and seasonal changes on survival and growth [40]. Unfortunately our time resolution (6 months) does not allow us to check for such correlation. We are currently developing an autonomous monitoring device that would enable us to drastically increase the monitoring frequency alongside the potential to collect environmental data such as sea surface temperature.

We were able to show the effect of sinking frames on coral mortality, yet we would like to include full substrate data into our regression analysis in the future. This will require the training of a new machine learning model to detect sedimentation and sand movement in monitoring pictures and generate frame-specific substrate data. We would also like to include current strength and turbidity, as it is suggested that it may have an impact on colony survival [41].

## 4 Methods

### 4.1 Coral transplantation

We transplant coral fragments on metal frames that are coated with resin and sand. These fragments are collected from wild and transplanted colonies that are found around our islands. They represent the local biodiversity of Acropora and Pocillopora genera. We collect a reasonable amount from each parent colony (maximum 20% of branches), that we fragment into pieces between 3cm and 8cm long.

Frames are assigned a reference number and physical tag. They are placed close to our islands, at depths ranging from 1 to 30 meters. We have limited our first study to a subgroup of our transplanted corals:

- Our coral frames are situated in the Maldives. The frames in this study are around Landaa Giraavaru (LG), Baa atoll (5.286ºN, 73.112ºE) and around Kuda Huraa (KH), North Male atoll (4.327ºN, 73.596ºE). Supplementary figure 1 shows their location.
- Of the 4 frame shapes that we are using (Small, Medium, Large and Heart), we are only analyzing the most common (Small). The smaller size enables the monitoring pictures to be more meaningful with each fragment representing a larger area in the picture. Each of these frames supports around 38 coral fragments.
- From 15 years of data, we are only using the frames that were transplanted after the 2016 mass bleaching event. They represent the largest volume of data points, as our program has consistently scaled up since its inception, and also the most consistent. Furthermore, the method we describe is not suitable for colonies older than 4-5 years. From that age they tend to merge with each other, and the metal frame is no longer visible, which prevents us from accurately identifying the individual colonies.

We also excluded 54 frames that have not been monitored for two years or more (usually as a result of a missing identification tag), and 5 frames with inconsistent monitoring pictures.

### 4.2 Monitoring

The monitoring process consists of capturing 4 pictures of each coral frame: front, left, back, and right, in this order, every six months. In this study, we are including 10,873 such monitoring sets, totaling 43,492 individual pictures. They are then automatically edited to maximize contrasts in red, green and blue channels. Finally, we normalize them using Contrast Limited Adaptive Histogram Equalization [42].

We automatically analyzed these pictures using the process presented in the next section.

### 4.3 Monitoring pictures analysis

#### 4.3.1 Coral detection

We have trained a convolutional neural network using 1103 manually annotated pictures which include 19890 annotations, representing 5 categories: Pocillopora, Acropora, Dead Coral, Bleached Coral and Frame Tag. We are using the Faster-RCNN network structure [22] with the ResNet 101 feature extractor [43], implemented with Tensorflow’s Object Detection library [44]. After training a model from scratch with these annotations, it is able to infer the position and category of objects in new pictures. We are provided with a classification, a bounding box and a score which represents the classification’s reliability.

#### 4.3.2 Structure detection

In order to reliably identify individual colonies, we need to know their position on the coral frame. Therefore, we need to detect the metal frame as it provides us with an objective reference in the 3D space.

To achieve this, we are using a U-Net [23] neural network, trained on 217 manually annotated pictures. The model outputs a binary mask of the area where frame bars were detected. It has a 97% accuracy (calculated using pixel-wise Softmax).

#### 4.3.3 Camera positioning

After detecting the metal structure in the picture, we need to find the 3D position of the camera relative to the frame. This is necessary to know the position of specific bars of the frame in the picture. To do this, we use stochastic optimization, which randomly simulates camera positions and returns the best match. We use differential evolution [46], implemented by the StochOPy package for Python [47].

#### 4.3.4 Merging data from the four angles

We then assign each detected fragment to a bar of the frame. The few fragments (3 to 6) that are on top of the frame are being ignored at this time, because their close proximity makes it impossible to accurately identify their supporting bar. Every fragment is assigned to a bar with a numeric position between 0 and 1 (left to right). We first record fragments from primary observations, that is from bars directly facing the camera. Then we make up for potential undetected fragments by merging fragments from secondary observations, if they were not already added in the first step. This enables us to merge data from all 4 monitoring pictures and have an outlook of what is on a frame at any specific monitoring date.

#### 4.3.5 Evolution tracking

To follow the evolution of each fragment, we go through the previously generated monitoring observations recursively. If the observation does not match with any previously observed fragment, it is considered a new fragment. To do this, we primarily check the position on the bar. If the position matches but there was a significant decrease in size (−20% at least), we assume the fragment has been replaced with a new one since the last monitoring event. Also, if a fragment was not detected in two consecutive monitoring sets, it is considered fallen.

We then consolidate that data. First, we fix the genus by taking the most frequent in all observations of that fragment. When a fragment was undetected / detected as dead in one monitoring set but reappears in the next set, we fix the mistake and declare it alive the whole time.

For each fragment, we set a transplantation date at which it is first detected. We also set a death date if death or fall is observed; we take the median date between the last live observation and the first dead / missing observation.

#### 4.3.6 Consistency check

We acknowledge that our method is prone to unfixable errors (due to the large number of observations and its automatic nature). We are using two different strategies to remedy this:

- Random checks: we select frames randomly to look at fragments that are incorrectly detected (fake positives, fake negatives, mis-attribution of observations to wrong fragments). This enables us to target the most frequent types of errors and adapt our method to reduce their frequency. This leads to manually annotating additional pictures and retraining of the models, or programmatic changes in our computer vision algorithms.
- Looking for outliers. Although our machine learning method is responsible for some mistakes, most significant errors derive from dataset inconsistencies. Usually, they arise because of two different frames that bear the same tag number, or improperly conducted monitoring (eg. going anticlockwise instead of clockwise). The best method we found to detect these issues is to look for outliers in our growth rate dataset. Indeed, inconsistent monitoring data tends to lead to different observations linked to the same fragment, which produces abnormal growth rates. Through manual checks we either confirmed the unlikely number, fixed the mistake when possible, or excluded the frame from the study.

We decided not to delete single fragments from the database so as not to introduce a bias. When mistakes were detected, as a last resort, we removed the whole frame from the study.

#### 4.4 Survival analysis

We are using the Kaplan-Meier model [27] to take right censoring into account in our results. Indeed we do not know the survival time of most of our fragments since they are still alive on the reef as we write this paper. It is a non-parametric model so it gives us the most realistic depiction of our survival results.

#### 4.4.1 Cox regression on the control group

In order to be sure that we are using a correct initial fragment size measurement, we are only considering fragments that were detected within three months following the frame transplantation date. This represents 9,759 fragments for the control group.

In order to identify the influence of depth, initial fragment size, genus and substrate, we used Cox’s multivariate regression [28], implemented in Python through the *lifelines* package [48]. We used Wei-Lin robust inference [49] which makes our results appropriate for practical use despite two of our covariates not verifying Cox’s proportional hazard hypothesis. The best result we obtained was when we considered interaction terms between genus and both depth and size.

The best-fitting formula we found is *C*(*Type*) *∗ log*(*Depth*)+ *C*(*Type*) *∗ Size* + *Sandy* : *Bottom*. It yields a 0.57 concordance and reasonable p-values. Table 2 shows the regression parameters for the control group.

**Table 1:** Number of manual annotations used for training for each category. The Average Precision is an indicator (0 to 1, 1 is best) of the model’s accuracy regarding each class [45].

| Category | Pocillopora | Acropora | Dead | Bleached | Tag | Total |
| --- | --- | --- | --- | --- | --- | --- |
| Annotations | 9547 | 6944 | 2165 | 804 | 430 | 19890 |
| Av. Precision | 0.89 | 0.76 | 0.72 | 0.73 | 0.68 | 0.76 |

**Table 2:** For each covariate, the coefficient is the natural logarithm of the hazard ratio (HR). HR low and HR high are the bounds of the 95% confidence interval for the hazard ratio.

|  | coef | HR | se(coef) | HR low | HR high | p |
| --- | --- | --- | --- | --- | --- | --- |
| **Type[Poc.]** | -0.87 | 0.42 | 0.11 | 0.34 | 0.52 | <0.005 |
| **log(Depth)** | -0.27 | 0.76 | 0.03 | 0.72 | 0.80 | <0.005 |
| **Type[Poc.]:log(Depth)** | 0.27 | 1.31 | 0.03 | 1.22 | 1.40 | <0.005 |
| **Size** | 0.06 | 1.06 | 0.02 | 1.02 | 1.10 | <0.005 |
| **Type[Poc.]:Size** | 0.05 | 1.05 | 0.02 | 1.01 | 1.09 | 0.02 |
| **Sandy:Bottom** | 0.25 | 1.28 | 0.03 | 1.20 | 1.36 | <0.005 |

#### 4.4.2 Cox regression with time-dependent variable

Again, we filter out the fragments that were detected more than 3 months after frame transplantation. This leaves 8,262 fragments for the House Reef (KH) and Water Villas (KH) zones.

As we added the time-dependant binary variable to account for environmental disruption, we cannot use the tradition Cox model and we had to use the derivative that is adapted to such variables [50]. The best-fitting formula we found is *C*(*Type*) *∗ Disrupted* + *C*(*Type*) : *Size*. Table 3 shows the hazard ratios with the associated 95% confidence intervals.

**Table 3:**
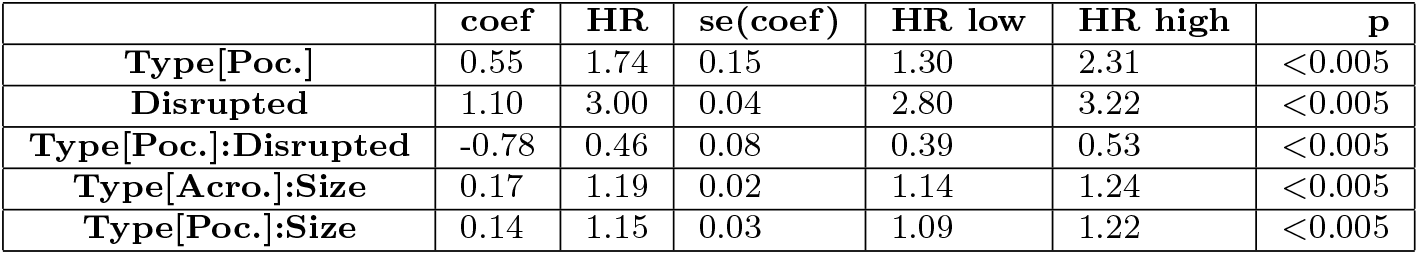
For each covariate, the coefficient is the natural logarithm of the hazard ratio (HR). HR low and HR high are the bounds of the 95% confidence interval for the hazard ratio.

#### 4.5 Size measurements

In order to check the correlation between real colony dimensions, and the approximations we get from machine learning annotations, we measured the dimensions of 150 coral colonies in our monitoring pictures. Their shape can be assimilated to a horizontal cylinder for smaller colonies (single branches), and a sphere for larger ones. In this dataset the tabling species are still small enough to be considered approximately spherical.

We consequently measured the apparent area of the colony, taking frame bar width as a size reference. In the case of spheres the apparent area is a circle, with the same radius as the sphere. For cylinders, the apparent area is a rectangle. Its height is twice the cylinder radius, and its width is the cylinder length.

##### 4.5.1 Linear size

As a linear length measurement, we are using the average between the height and length of the bounding box. To assess its correlation to real world dimensions, we compared it to a representative length derived from the previously mentioned measurements: the radius for spheres, and *radius*^2*/*3^*· length*^1*/*3^ for cylinders.

We observe a linear correlation (*R*^2^ = 73%) between the machine learning calculated measurement (*L*_*calculated*_ = 0.5*·* (*height* + *width*)) and the manually measured dimension:

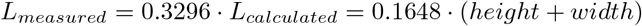

The residuals have an average of −0.022mm and a standard deviation of 0.80cm. Visual analysis suggests that they are randomly distributed (see figure 7a). We then consistently used this formula to calculate coral growth.

**Figure 7:**
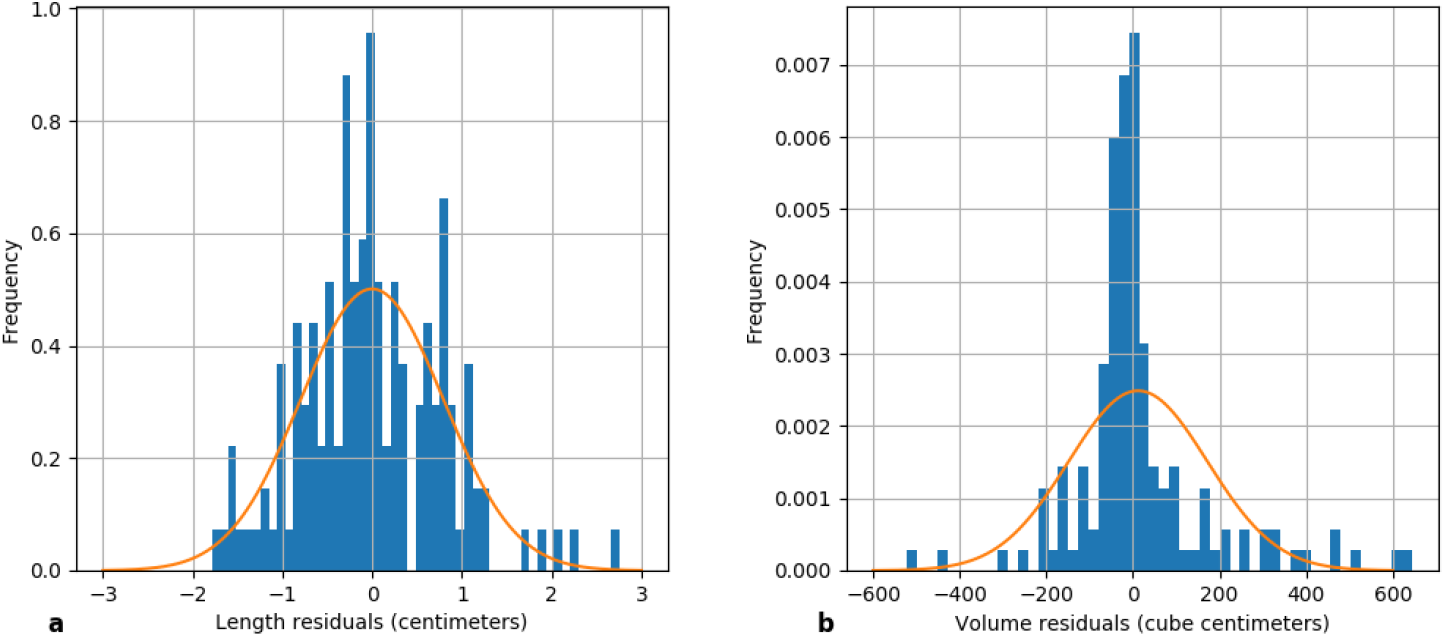
Distribution of length (**a**) and volume (**b**) residuals. We show in orange the gaussian distribution of the same average and standard deviation as the residuals.

##### 4.5.2 Volume

Diameter of coral colonies was shown to be a good indicator of their shelter volume [30]. We need to check our machine learning estimates against the volume that we measured manually from the pictures. We converted the manually measured surface area to volume by multiplying it by the right length: 4*/*3 *radius* for a sphere and *π/*2*· radius* for a cylinder. Then we compared it to the estimate that we get from the bounding boxes dimensions generated by automatic annotations: (*height · width*)^1^.^5^. This gave us the following relationship with *R*^2^ = 72% :

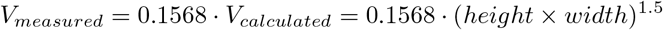

The residuals have a mean of 0.012 liters and a standard deviation of 0.160 liters. Visual analysis suggests that they are randomly distributed (see figure 7b). We then consistently used this formula to automatically infer the volume of each colony.

## Data and code availability

All the code that we wrote for this article, as well as the list of dependencies we used, are available on GitHub under the GNU GPL v3 license [51].

Our frame and fragment database is available on SEANOE under the Creative Commons BY-SA license [25].

Our photographic database is available on our website [52].

## Competing interests statement

The authors declare the following competing interests: Gaé tan Morand and Simon Dixon are employed by Reefscapers. Thomas Le Berre is managing director of Reefscapers.

## Acknowledgements

We thank all the marine biologists who worked on this restoration program and made this study possible by placing and maintaining thousands of coral frames.

We thank Four Seasons Resorts Maldives for their continuous support since the beginning of this restoration program and all the generous guests and online sponsors who helped finance coral transplantation.

We thank Hughes Talbot for his useful advice regarding frame structure detection, and Laurent Goetz for his precious comments and improvements to the manuscript.

## Notes

https://doi.org/10.17882/77361

### Competing Interest Statement

Gaetan Morand and Simon Dixon are employed by Reefscapers. Thomas Le Berre is managing director of Reefscapers.

