## Supplementary Figure 1 for "Assessment of coral restoration’s contribution to reef recovery using machine learning"

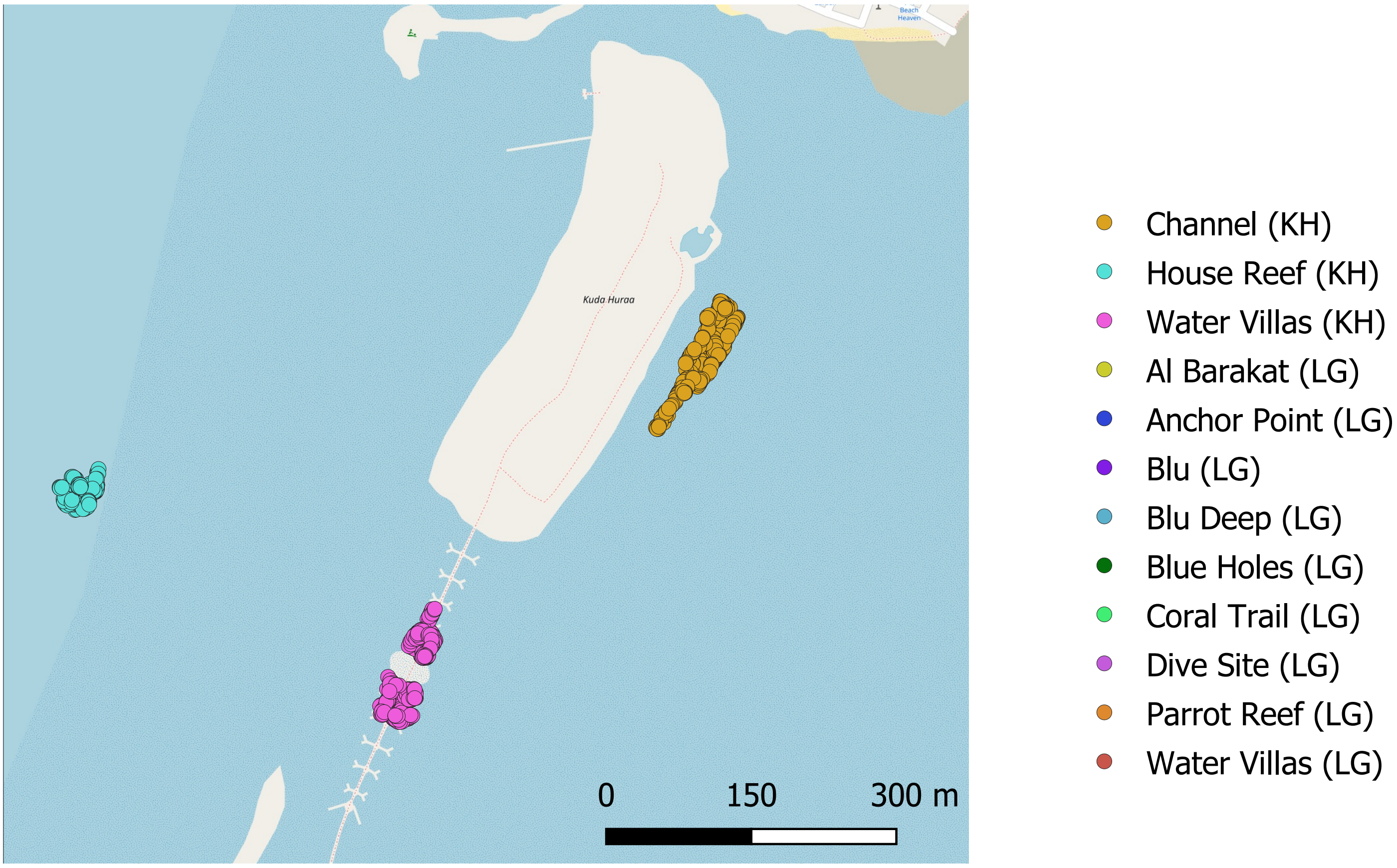


Kuda Huraa

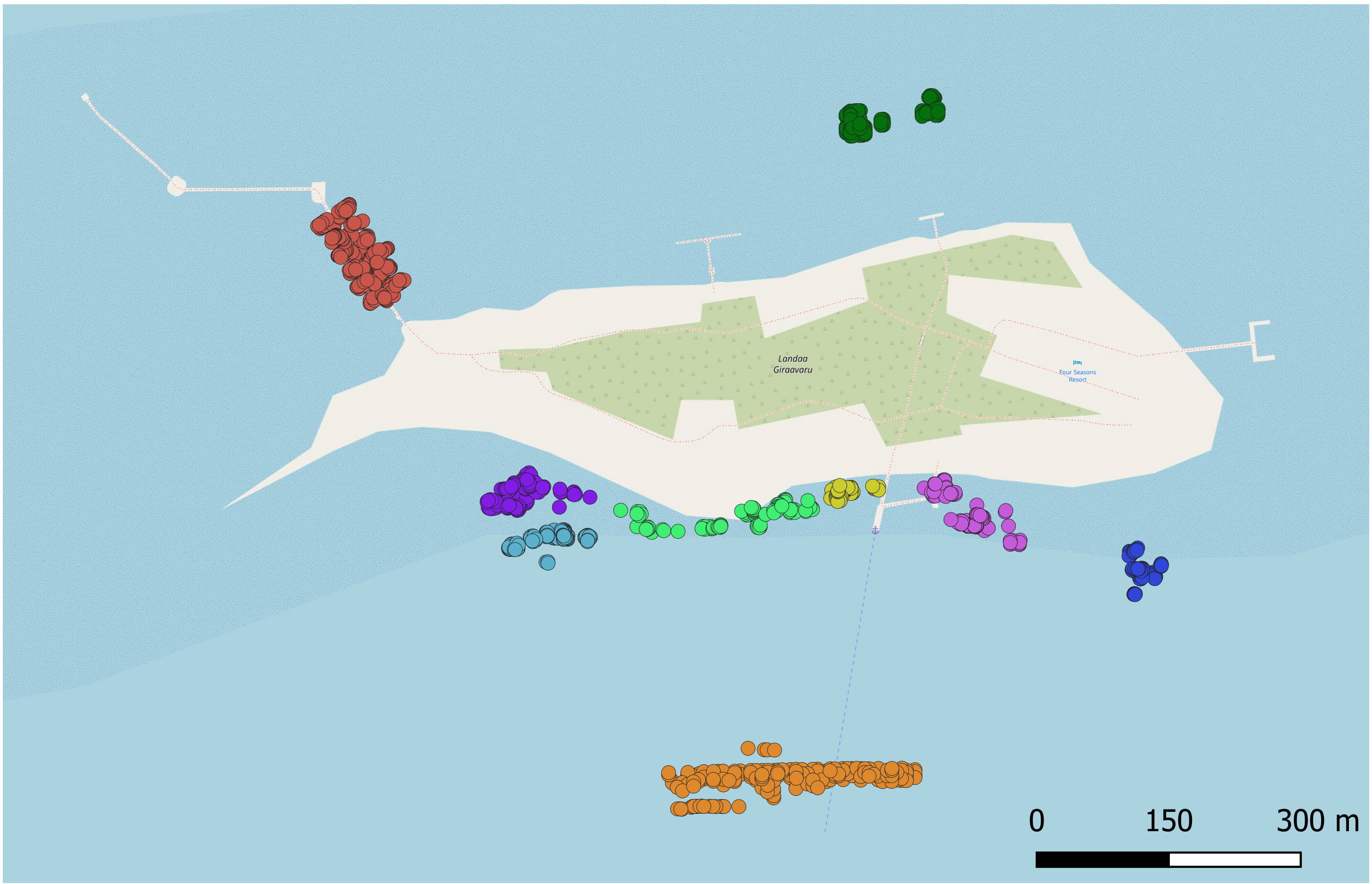


Landaa Giraavaru

Supplementary Figure 1: Location of the 2,236 frames in the 12 major geographical zones. Kuda Huraa and Landaa Giraavaru are 119 km apart.

Base maps and data from OpenStreetMap and OpenStreetMap Foundation
